# Identification of functional genes underlining growth in large yellow croaker

**DOI:** 10.1101/2023.11.21.567991

**Authors:** Jing Wen, Zhufeng Yue, Jiaojun Ji, Yingbo Yuan, Meng Zhou, Bo Zhang, Yongjie Zhang, Kun Ye, Zhiyong Wang, Dan Jiang, Ming Fang

## Abstract

Large yellow croaker (*Larimichthys crocea*) is the largest cultured marine fish in China, although its economic benefits are significant, its growth cycle is long and it takes 2 years to market. Identification of genomic variations and genes affecting the growth of *L. crocea* is of great significance for breeding fast-growing strains by molecular means. We utilized whole genome resequencing to identify a highly effective GWAS locus on chromosome 6 of the genome, and identified some candidate genes that affect growth. This study provides a theoretical basis for analyzing the genetic mechanism of growth traits of *L. crocea* and cultivating fast-growing varieties.

## Introduction

Large yellow croaker (*Larimichthys crocea*) is a type of warmly-clustered migratory fish, belongs to the order Perciformes, family Sciaenidae, genus *Larimichthys*. It is the largest cultured marine fish in China, with a production of 25.6 million tons in 2022. The growth rate of large yellow croaker is slow, and it takes 2 years to grow to the market specification. The long breeding cycle leads to a sharp increase in breeding risks such as diseases and typhoons. Cultivating new varieties of fast-growing yellow croaker is of great significance for improving the economic benefits of yellow croaker farming.

Breeding is an important way to improve the economic benefits of large yellow croaker farming. The individual growth rate of large yellow croaker varies greatly, and there is great room for breeding. Molecular breeding is an important means of accelerating genetic selection. Mining molecular markers from a molecular perspective is of great significance for improving the growth rate of *L. crocea*. Genome-wide association has been widely used in the identification of major genes in fish. Yu et al. used the ddRAD-seq method to locate the whole genome of quantitative trait loci (QTLs) related to the growth of *Epinephelus coioides*, 17 candidate functional genes possibly related to the growth of *Epinephelus coioides* were detected ^[1]^; Liu et al. conducted genome-wide QTL analysis on the growth traits of tilapia and detected a major QTL controlling growth traits at the LG1, a candidate gene *ghr2* in this QTL was significantly correlated with the weight of tilapia, explaining 13.1% of phenotypic variation. Real time fluorescence quantitative PCR results showed that different genotypes of *ghr2* affected the expression level of the *igf1* gene, indicating that *ghr2* may be an important gene underling growth of tilapia ^[2]^. In large yellow croaker, Dong et al identified a number of candidate genes underling growth of large yellow croaker with RAD-seq technology ^[1]^.

The effectiveness of GWAS also depends on the density of genomic SNPs, compared with simplified genome sequencing and SNP chips, whole genome resequencing can obtain a higher density of genetic variation, which can improve the efficiency of GWAS. In this study, we will study the major genes of growth traits of *L. crocea* in the whole genome by means of high-throughput whole genome resequencing. We conducted the association study on the growth traits of juvenile and adult yellow croaker, and found that the two growth stages were controlled by the same main genome region. This study provides molecular markers for the breeding of fast growing traits in *L. crocea*.

## Materials and methods

### Sample collection

*Adult fish sample:* We randomly selected ∼1000 individuals with different growth rates from 2-year-old yellow croakers cultured in the same cage at the Sandu Ao Yellow Croaker Breeding Base in Ningde City, Fujian Province in December 2019. Before sample collection, 674 individuals with extreme growth phenotypes were domesticated indoors at the breeding base for two weeks, and their body weight (BW) was measured Phenotypes of growth related traits such as Body Length (BL) and Body Height (BH). The average body length is 390.3 ± 154.1 g, with an average body length of 26.1 ± 3.1 cm. The tail fins were collected and DNA was extracted. We complied the requirements of the Animal Ethics Committee of the School of Fisheries at Jimei University for experimental animal operation.

*Young fish sample:* We collected a total of 500 young yellow croakers in the same cage at the Guanjingyang Yellow Croaker Breeding Base in Sandu Ao, Ningde City, Fujian Province in August 2022. The average body length of the fry was 7.27±0.86cm, body weight was 7.15±2.6g, and the tail fins were collected for DNA extraction.

### Sequencing data processing

We used fastqc software (https://www.bioinformatics.babraham.ac.uk/projects/fastqc/) for quality control of the clean data, used BWA-MEM software (http://bio-bwa.sourceforge.net/) ^[3]^ to align the high-quality sequencing reads to the reference genome of large yellow croaker (both the reference genome file and annotation file are assembled in our laboratory); we used Samtools software ^[4]^ to convert the sam files into bam files, and then sorted and created an index using Sambamba software (https://github.com/4biod/sambamba/releases). The markdup tool in ^[5]^ was used to remove duplicate fragments generated during the PCR amplification step of library preparation, and then GATK v4.1.9.0 (Genome Analysis Toolkit) software is used (https://gatk.broadinstitute.org/hc/en-us) with HaplotypeCaller, CombineGVCFs, and GenotypeGVCFs modules ^[6]^ for discovering SNP and small InDel variants. The SNPs and InDels were filtered separately and finally merged together. We filtered the SNPs with the parameters “QD<2.0 | | MQ<40.0 | | FS>60.0 | SOR>3.0 | MQRankSum<-12.5 | | ReadPosRankSum<-8.0”, and filtered InDels with parameters “QD<2.0 | | FS>200.0 | SOR>10.0 | MQRankSum<-12.5 | | ReadPosRankSum<-8.0”. Finally, we used Beagle v4.1 software ^[7]^ t impute missing variants.

After genotyping all individual samples, we used bcftools v1.11 ^[8]^ to merge the VCF files that record genotype information for all sample individuals. The merged VCF file contains all SNP genotype information generated by whole genome resequencing of 674 sample individuals, and then used Plink v1.9 software ^[9]^ to format SNP genotype information for the genome-wide association analysis.

### Genome-wide association study

This study used EMMAX software for genome-wide association study, which has the advantages of simple operation and fast calculation speed. By reducing the dimensionality of the calculation function, the software can complete millions of SNPs in a very short time. Therefore, this software is well suited for whole genome association analysis of large sample populations. The EMMAX software uses a mixed linear model to perform correlation analysis on phenotype, genotype, and phylogenetic data. The statistical model formula is as follows (1):

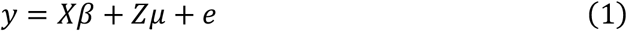

y is a vector used to represent the phenotypic values of the studied trait, and X is the corresponding fixed effect correlation matrix, β Is the vector corresponding to the fixed effect, μ, is the vector corresponding to individual additive genetic random effects, Z is the random effect correlation matrix, that is, the effect matrix of SNPs (the genotypes “AA”, “Aa”, and “aa” of SNPs correspond to “0”, “1”, and “2” respectively), and e is the vector composed of all random errors. This software uses the Bonferroni correction method to correct the p-value. The EMMAX software ultimately conducts correlation analysis between SNP and trait phenotype through the EMMAX package.

Using the Cmplot ^[10]^ package in R language to draw Manhattan plot, the threshold p-value for significant association between SNP and the white spot disease resistance traits of large yellow croaker is 0.05/N, and the extremely significant threshold p-value is 0.01/N (N represents the number of effective SNPs in the dataset). Finally, genes in the 1 Mb region upstream and downstream of the significantly associated SNP loci were annotated using NCBI and Genecard related gene function annotation websites to determine whether their functions are related to the resistance to surface white spot disease in yellow croaker. Finally, candidate functional genes related to the resistance to surface white spot disease in yellow croaker were identified.

## Results

### Adult fish results

A total of 674 large yellow croaker DNA samples were re-sequenced using the Illumina high-throughput sequencing platform, resulting in an average of 14,783,875 raw reads, 14,608,918 clean reads, 4.44 GB raw data, 4.38 GB clean data. The average GC content is 42.11%. Using GATK software, we conducted SNP site mining on the whole genome re-sequencing data of 674 large yellow croaker DNA samples, and a total of 30,214,092 SNPs were discovered for genome-wide association analysis.

The EMMAX was used to conduct genome-wide association analysis on 30,214,092 SNPs from 674 whole genome re-sequencing data of *L. crocea* and 3 growth related traits, including body weight, body length and body height are studied. The significant threshold of the association *p*-value was 1.655E-09 (0.05/30,214,092), The threshold for highly significant association is *p*-value=3.310E-10 (0.01/30,214,092). It can be seen that the GWAS profiles for the three growth traits are very similar, a significant SNP cluster was formed on chromosome 6 (Figure 1). The most significant SNP was chr6_19106799, the minor allelic frequency (MAF) was 0.3093. We list the detailed information of all significant SNPs in Table 1. Based on the gene sequences in the 1 Mb region upstream and downstream of the lead variant of chr6 GWAS locus, we annotated the primary functional genes with uniport and NR database. The annotated functional genes are listed in Table, including *Ccnl1, Veph1, Pdcd10b, Tipar* and *Serpini1* and so on.

**Table 1.**
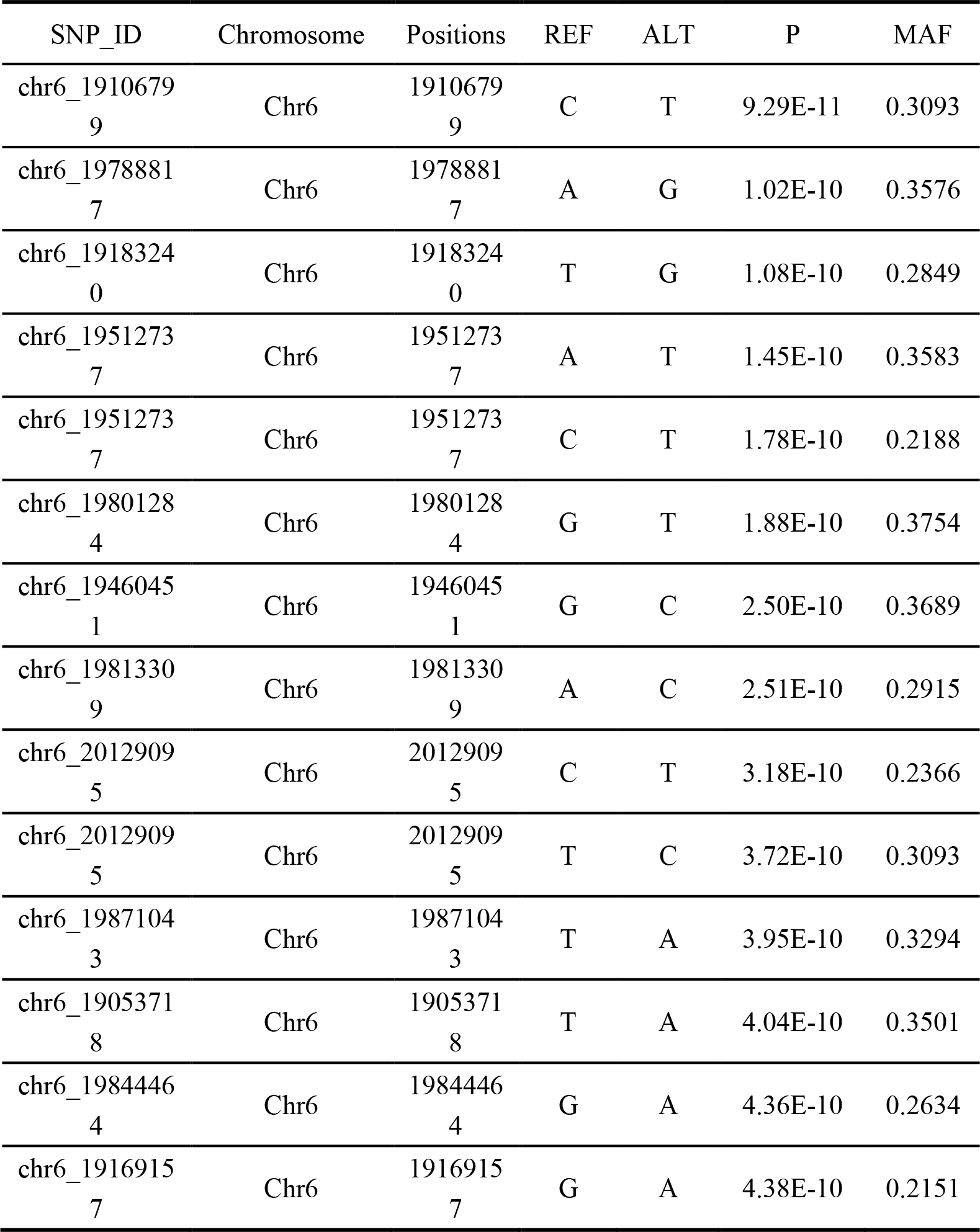
Significant SNPs of the GWAS of body weight of adult fish.

| SNP_ID | Chromosome | Positions | REF | ALT | P | MAF |
| --- | --- | --- | --- | --- | --- | --- |
| chr6_1910679<br>9 | Chr6 | 1910679<br>9 | C | T | 9.29E-11 | 0.3093 |
| chr6_1978881<br>7 | Chr6 | 1978881<br>7 | A | G | 1.02E-10 | 0.3576 |
| chr6_1918324<br>0 | Chr6 | 1918324<br>0 | T | G | 1.08E-10 | 0.2849 |
| chr6_1951273<br>7 | Chr6 | 1951273<br>7 | A | T | 1.45E-10 | 0.3583 |
| chr6_1951273<br>7 | Chr6 | 1951273<br>7 | C | T | 1.78E-10 | 0.2188 |
| chr6_1980128<br>4 | Chr6 | 1980128<br>4 | G | T | 1.88E-10 | 0.3754 |
| chr6_1946045<br>1 | Chr6 | 1946045<br>1 | G | C | 2.50E-10 | 0.3689 |
| chr6_1981330<br>9 | Chr6 | 1981330<br>9 | A | C | 2.51E-10 | 0.2915 |
| chr6_2012909<br>5 | Chr6 | 2012909<br>5 | C | T | 3.18E-10 | 0.2366 |
| chr6_2012909<br>5 | Chr6 | 2012909<br>5 | T | C | 3.72E-10 | 0.3093 |
| chr6_1987104<br>3 | Chr6 | 1987104<br>3 | T | A | 3.95E-10 | 0.3294 |
| chr6_1905371<br>8 | Chr6 | 1905371<br>8 | T | A | 4.04E-10 | 0.3501 |
| chr6_1984446<br>4 | Chr6 | 1984446<br>4 | G | A | 4.36E-10 | 0.2634 |
| chr6_1916915<br>7 | Chr6 | 1916915<br>7 | G | A | 4.38E-10 | 0.2151 |

**Table 2.** Functional genes near significant SNPs of adult fish.

| SNP_ID | Chromosome | Region | Genes |
| --- | --- | --- | --- |
| chr6_19106799 | Chr6 | Intron | *Znf429* |
| chr6_19788817 | Chr6 | Intergenic | - |
| chr6_19183240 | Chr6 | Intron | *Scn2b* |
| chr6_19512737 | Chr6 | Intron | *Si* |
| chr6_19512737 | Chr6 | Intergenic | *Slitrk3*,<br>*Bche* |
| chr6_19801284 | Chr6 | Intergenic | - |
| chr6_19460451 | Chr6 | Intergenic | - |
| chr6_19813309 | Chr6 | Intergenic | - |
| chr6_20129095 | Chr6 | Intron | *Ccn1l*,<br>*Veph1*,<br>*Pdcd10b*,<br>*Tipar*,<br>*Serpini1* |
| chr6_19906254 | Chr6 | Intergenic | *Serpini1*,<br>*Golim4* ,<br>*Slc66AA1*,<br>*Melt* |
| chr6_19053718 | Chr6 | Intergenic | *Znf429*,<br>*Megf6* |
| chr6_19844464 | Chr6 | Intergenic | - |
| chr6_19169157 | Chr6 | Intergenic | *Wdr49*,<br>*Zbbx* |

**Figure 1.**
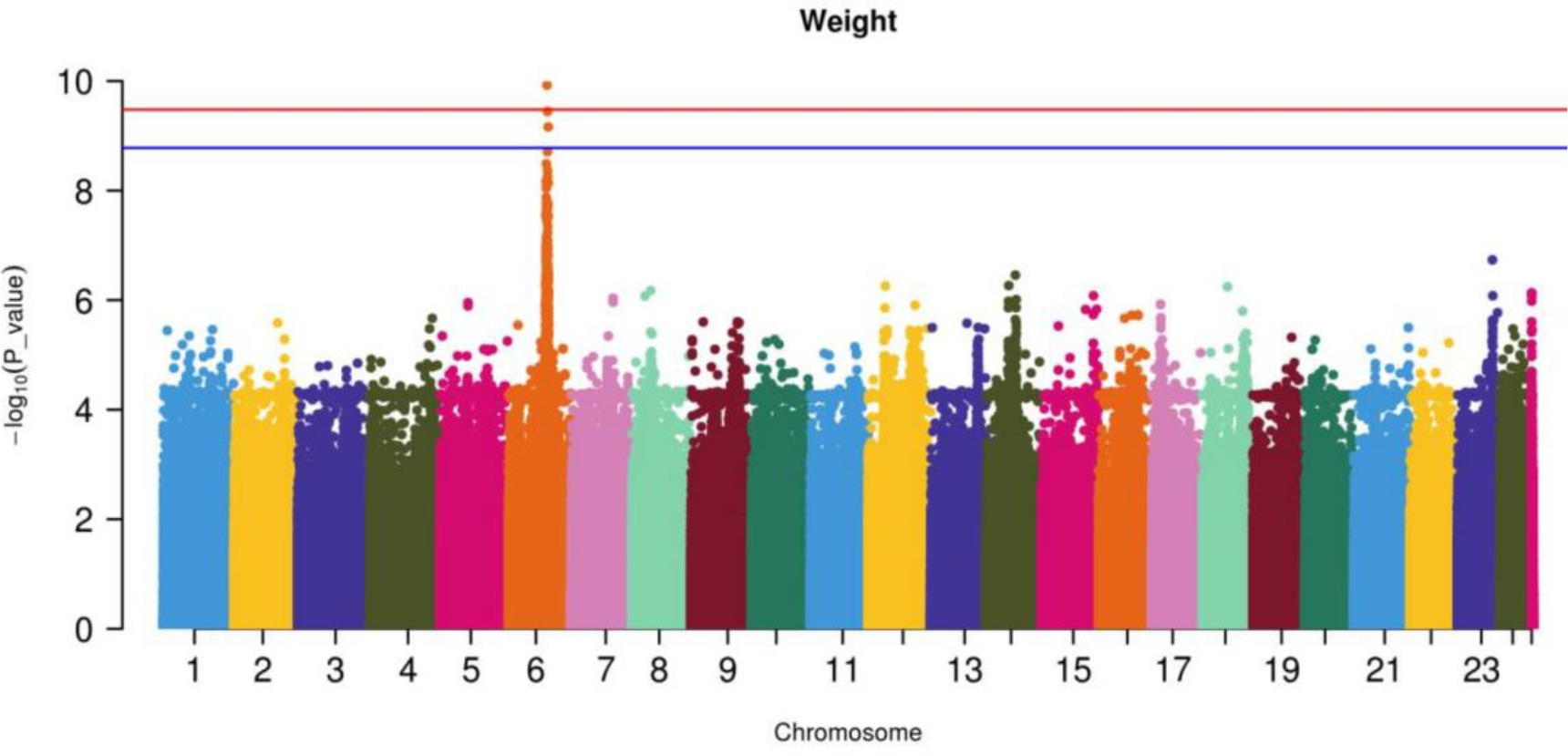
GWAS results of the body weight of adult fish.

### Juvenile fish results

Totally, 5823030 variants were discovered with whole genome resequencing of 500 juvenile fish, the GWAS analysis was conducted with EMMAX. We declared the significance of GWAS loci at genome-wide level with thresholds 1.717319e-09 (0.01/5823030) and 8.586595e-09 (0.05/5823030), we show the results of body weight and length in Figure 2 and 3, four significant GWAS loci on chromosomes 6, 17 and 19 were identified for body weight, respectively, ande four clusters were identified on chromosome 6, 19 and 20. The most significant GWAS locus for both body weight and length was on Chr6, the lead variant for body weight GWAS locus was at 20,247,596 bp with p-value 3.76E-09 and allele frequency 0.112. The position of the lead variant is very close to that of the adult fish chr6_19106799, suggesting both adult fish and juvenile fish are potentially share the same GWAS loci. We list the detailed information of significant SNPs in Table 3. We also annotated the functional genes of chr6 in Table 4, among which, several functional genes such as *Ccnl1, Zbbx, Wdr49, Tiparp, Veph1* and *Pdcd10b*, are overlapping with genes identified in adult fish, implying the importance of these genes on growth of *L. crocea*.

**Table 3.** Significant SNPs of the GWAS of body weight of juvenile fish.

| SNP_ID | Chromosome | Positions | REF | ALT | P | MAF |
| --- | --- | --- | --- | --- | --- | --- |
| chr6_2024759<br>6 | Chr6 | 2024759<br>6 | A | C | 3.76E-09 | 0.112 |
| chr6_2010613<br>8 | Chr6 | 2010613<br>8 | C | T | 8.21E-09 | 0.126 |
| chr6_2010614<br>8 | Chr6 | 2010614<br>8 | T | C | 8.21E-09 | 0.126 |
| chr6_2014812<br>5 | Chr6 | 2014812<br>5 | G | A | 1.84E-08 | 0.147 |
| chr6_1991173<br>3 | Chr6 | 1991173<br>3 | C | T | 1.90E-08 | 0.121 |
| chr6_1976243<br>7 | Chr6 | 1976243<br>7 | G | T | 3.08E-08 | 0.122 |
| chr6_1991436<br>8 | Chr6 | 1991436<br>8 | A | G | 3.61E-08 | 0.122 |
| chr6_2045240<br>2 | Chr6 | 2045240<br>2 | G | A | 3.69E-08 | 0.089 |
| chr6_2027599<br>3 | Chr6 | 2027599<br>3 | A | G | 3.82E-08 | 0.139 |
| chr6_2185592<br>9 | Chr6 | 2185592<br>9 | C | A | 4.16E-08 | 0.283 |
| chr6_2012913<br>6 | Chr6 | 2012913<br>6 | T | C | 4.52E-08 | 0.143 |
| chr6_1991450<br>3 | Chr6 | 1991450<br>3 | A | T | 4.54E-08 | 0.070 |
| chr6_2032255<br>5 | Chr6 | 2032255<br>5 | T | G | 4.78E-08 | 0.112 |
| chr6_1993662<br>3 | Chr6 | 1993662<br>3 | G | A | 4.87E-08 | 0.091 |

**Table 4.**
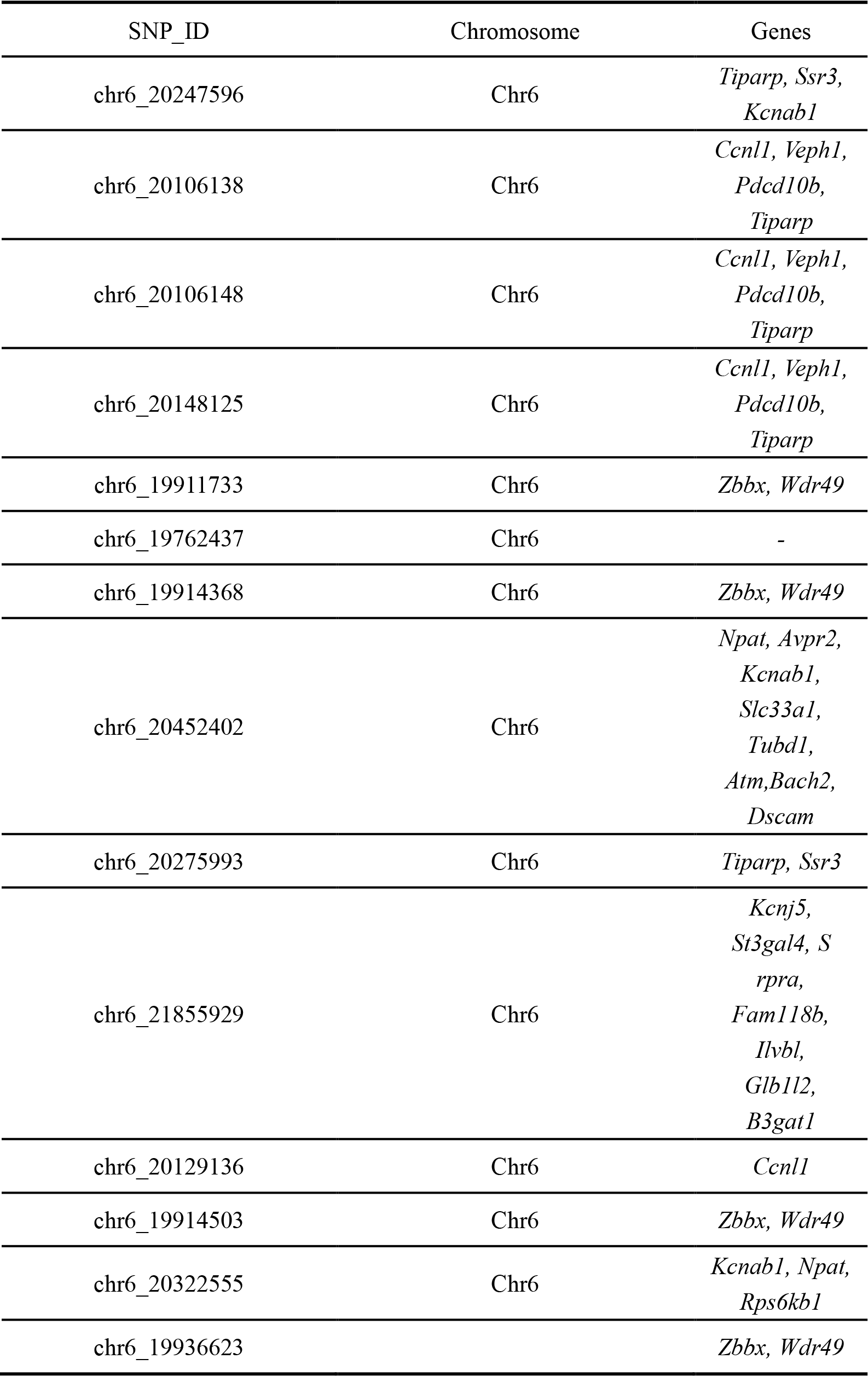
Functional genes near significant SNPs of juvenile fish.

**Figure 2.**
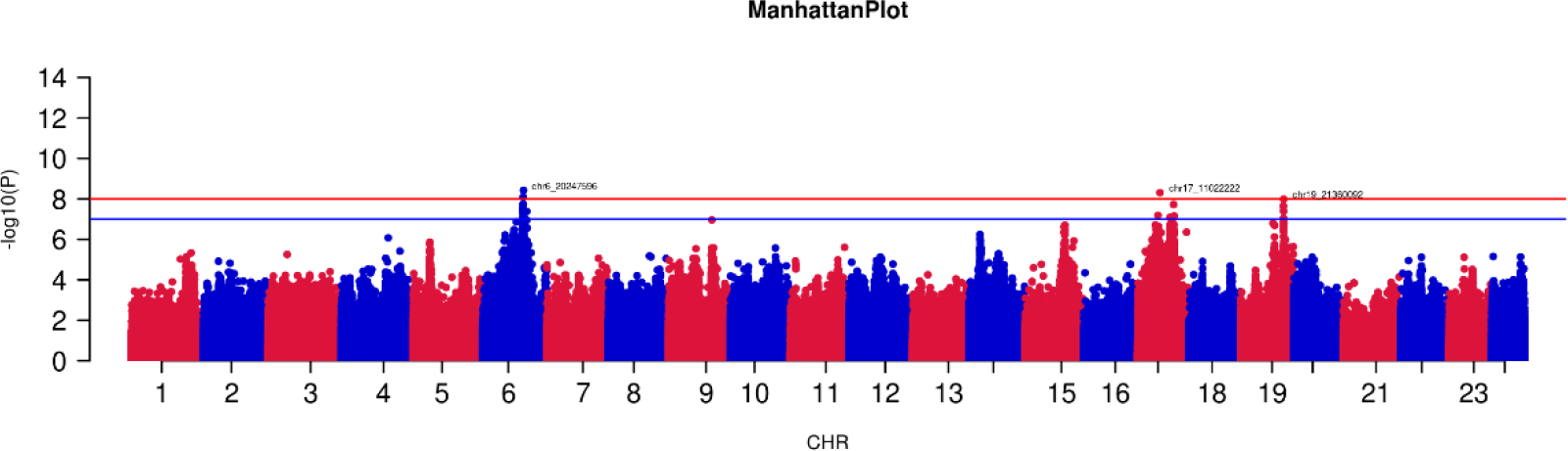
GWAS results of the body weight of adult fish.

**Figure 3.**
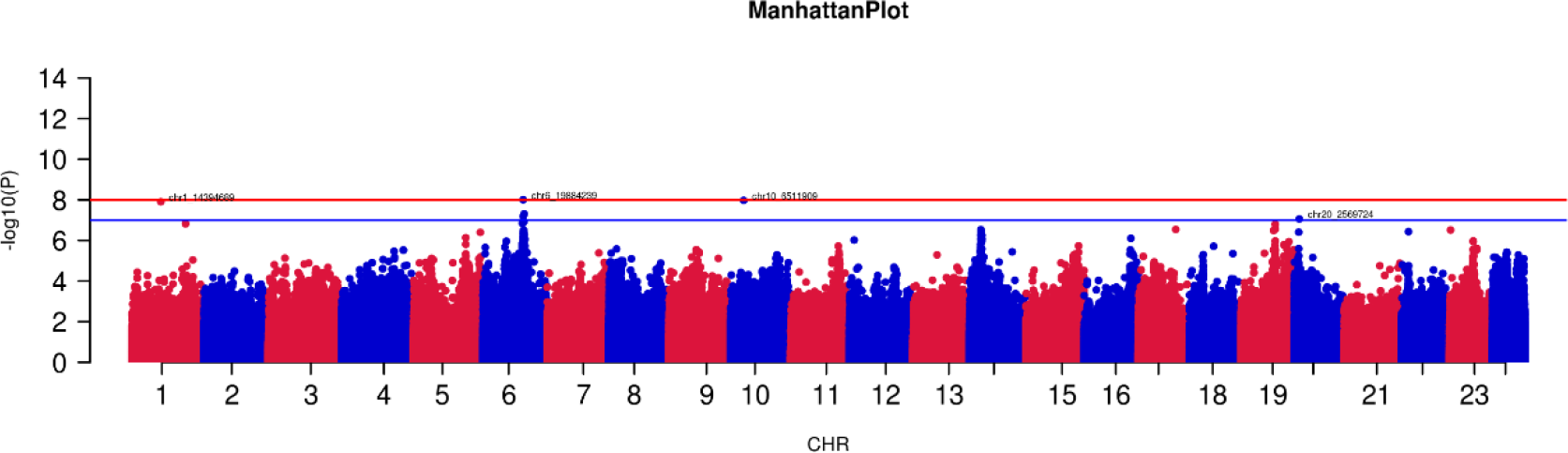
GWAS results of the body length of adult fish.

## Discussions

The results showed that both adult and juvenile fish detected a common locus on Chr6, indicating a similar genetic basis of two growth stage, the deep study of the GWAS locus will be meaningful for molecular selection.

We identified several functional important candidate genes, but we still have no enough evidence to confirm they are causative. These genes have not been reported to be directly related to the growth of other species, which increases the difficulty to narrow down the candidate genes. Now, we have employed gene editing and transgenes techniques to validate several genes including *Si, Slitrk3, Ccnl1, Veph1, Pdcd10b, Tiparp* and *Golim4*, and we will discover the causative genes in future.

We use whole genome re-sequencing to identify the genetic variation that leads to the growth of *L. crocea*. The whole genome re-sequencing can obtain rich genetic variation, including not only all potential SNP genetic variations, but also Indel variations, which has the possibility of detecting causal mutations. However, the existence of linkage imbalance makes it difficult to identify causal mutations from a large number of variants that are in linkage. Nevertheless, identifying causal mutations is our next task.

## References

1. Yu H, You X X, Li J, et al. Genome-wide mapping of growth-related quantitative trait loci in Orange-spotted grouper (Epinephelus coioides) using double digest restriction-site associated DNA sequencing (ddRADseq)[J]. International Journal of Molecular Sciences, 2016, 17(4): 501.

2. Liu F, Sun F, Xia J H, et al. A genome scan revealed significant associations of growth traits with a major QTL and GHR2 in tilapia[J]. Scientific Reports, 2014, 4: 7256.

3. Li H. Aligning sequence reads, clone sequences and assembly contigs with BWA-MEM[J]. arXiv preprint arXiv: 1303.3997, 2013.

4. Li H, Handsaker B, Wysoker A, et al. The sequence alignment/map format and SAMtools[J]. Bioinformatics, 2009, 25(16):2078–2079.

5. Tarasov A, Vilella A J, Cuppen E, et al. Sambamba: fast processing of NGS alignment formats[J]. Bioinformatics, 2015, 31(12): 2032–2034.

6. Mckenna A, Hanna M, Banks E, et al. The genome analysis toolkit: a MapReduce framework for analyzing next-generation DNA sequencing data[J]. Genome research, 2010, 20(9): 1297–1303.

7. Ayres D L, Darling A, Zwickl D J, et al. BEAGLE: an application programming interface and high-performance computing library for statistical phylogenetics[J]. Systematic Biology, 2012, 61(1): 170–173.

8. Danecek P, McCarthy S A. BCFtools/csq: haplotype-aware variant consequences. Bioinformatics, 2017, 33(13): 2037–2039.

9. Purcell S, Neale B, Todd-Brown K, et al. PLINK: a tool set for whole-genome association and population-based linkage analyses[J]. American Journal of Human Genetics, 2007, 81(3): 559–575.

10. Tan Q S, Wang F, Xie S Q, et al. Effect of high dietary starch levels on the growth performance, blood chemistry and body composition of gibel carp (Carassius auratus var. gibelio). Aquaculture Research, 2009, 40(9): 1011–1018.

